# Molecular mechanism of coilin interaction with core snRNPs

**DOI:** 10.64898/2026.03.10.710772

**Authors:** Nenad Radivojević, Veronika Holotová, Martina Groušlová, Utz Fischer, David Staněk

## Abstract

Cajal bodies (CBs) are nuclear membrane-less organelles that accumulate various short non-coding RNAs and facilitate their biogenesis. They also function in quality control during the assembly of small nuclear ribonucleoprotein particles (snRNPs), sequestering immature or defective complexes. In this paper, we show that coilin, the key scaffolding protein of CBs, is the factor that discriminates between mature and immature snRNPs. We provide evidence that the C-terminus of coilin contains a bipartite snRNP-interaction module composed of a nonspecific RNA-binding region formed by RG repeats and a Tudor-like domain that interacts specifically with Sm proteins. The Tudor-like domain contains two conserved loops that protrude from the core barrel and bind the Sm proteins E, F, and G. Both the RNA-binding and Sm-binding regions are essential for productive interactions between coilin and core snRNPs, providing a molecular explanation for the specificity of coilin-mediated sequestration of immature snRNPs in Cajal bodies.

## Introduction

RNA splicing is a key step in mRNA maturation in virtually all eukaryotes. This process is catalyzed by the spliceosome, a highly dynamic multi-megadalton complex composed of five small nuclear ribonucleoprotein particles (snRNPs) and numerous auxiliary factors. snRNPs are multicomponent complexes containing a non-coding small nuclear RNA (snRNA), a heptameric Sm or Like-Sm (LSm) ring, and a set of snRNP-specific proteins. Except for U6 snRNA, U1, U2, U4 and U5 snRNAs are transcribed by RNA polymerase II and immediately after transcription exported to the cytoplasm by a PHAX containing cap-binding complex ^1–3^. In the cytoplasm, snRNAs interact with the survival of motor neuron (SMN) complex, which re-structures pre-snRNAs and facilitates the loading of the Sm proteins onto the Sm binding site, forming the core snRNP complex ^4–6^. The 5’ cap of the snRNA is then hypermethylated and, together with the assembled Sm proteins, serves as a signal for re-import of the core snRNP into the nucleus ^7–11^.

In the nucleus, core snRNPs first appear in CBs, where snRNAs undergo ribose methylation and pseudouridylation, while their 3’ ends are trimmed to produce mature transcripts ^12–15^. CB also serves as a hub for the final maturation and recycling of splicing-competent U2 and U4/U6•U5 tri-snRNPs ^16–22^. The imported core snRNPs are converted into mature (i.e. splicing competent) particles in a step-wise manner. The U2 core snRNP first associates with SNRPA1(U2A’)/SNRPB2(U2B’’) dimer, forming an early 12S assembly intermediate ^23^. SF3B and SF3A complexes are subsequently added, generating the 15S and mature 17S particles, respectively ^23–25^. Similarly, the tri-snRNP is assembled sequentially. Formation of the U4/U6 di-snRNP is followed by the addition of the U5 snRNP, which itself is assembled in several steps ^21,26–29^.

snRNP biogenesis involves several quality control checkpoints that monitor their correct maturation. First, the formation of the snRNA export complex is monitored in CBs ^30^. In the cytoplasm, Sm protein production is tightly controlled ^31,32^, the presence of the 5’ cap and the Sm binding site is scrutinized by GEMIN5 ^33,34^, and snRNAs that fail to acquire the Sm ring are rapidly degraded in the cytoplasm ^35–39^. In addition, processing of the 3’-end of the pre-snRNA neglects unstable snRNA variants, leaving them vulnerable to exosome degradation, providing an additional layer of quality control ^40,41^. Finally, core snRNPs that fail to form the splicing-competent complexes are sequestered in CBs ^16,17,42^. Targeting and sequestration of immature snRNPs in CBs requires the presence of the Sm ring, but the CB factor(s) that interact with the core snRNP and discriminate between immature and mature complexes are unknown ^43^.

Coilin is the key scaffolding protein of CBs, and its knockout results in CB disintegration across all studied organisms and cell types ^44–49^. Protein-wide analyses identified multiple coilin interactors, including ribosomal proteins and components of snoRNP, snRNP and SMN complexes (^50–52^ for review see ^53^). Although it has been suggested that coilin enhances and coordinates the metabolic pathways of various non-coding RNAs and RNPs, its precise role remains unclear ^51,52,54–59^. Human coilin has 576 amino acids, but only ∼80 amino acids at the N-terminus and ∼140 amino acids at the C-terminus are conserved and structured ^60,61^. The N-terminus contains a self-interaction domain, which is essential for CB formation ^60,62^. The central part of the protein is intrinsically disordered and, apart from the two nuclear localization signals adjacent to the N-terminus, harbors no apparent functional domains ^53,63^. The C-terminal region contains an arginine/glycine-rich domain called the RG box, followed by the Tudor-like domain found at the very C-terminus ^61^. This region has been reported to interact with snRNPs and Sm proteins, and this interaction is essential for CB formation ^57,58,63,64^. Arginines in the RG box are symmetrically methylated and serve as a binding platform for the SMN protein by binding to its central Tudor domain ^58,65,66^. A canonical Tudor domain adopts a β-barrel structure formed by four antiparallel β-strands connected by short loops. Four conserved aromatic residues within two of these loops (β1-β2 and β3-β4) form an aromatic cage that binds methylated lysine and arginine residues with high affinity and specificity ^67,68^. Unlike canonical Tudor domains, the Tudor-like domain of coilin lacks the characteristic aromatic cage. While the core β-barrel structure is preserved, β1-β2 and β3-β4 loops are extended and flexible, displacing the aromatic residues required for the binding of methylated amino acids ^61^. These atypical loops are conserved among coilin homologs but are absent in canonical Tudor domains, suggesting a divergent function or interaction specificity.

In this study, we performed a detailed molecular characterization of coilin interaction with snRNPs to test the hypothesis that coilin is the missing factor that sequesters immature snRNPs in CBs. We identified the RG box as a thus far unknown RNA-binding domain and the conserved loops protruding from the Tudor-like core as key mediators of interactions with Sm proteins. Our data suggest that the RG box and the Tudor-like domain constitute a bipartite interaction platform that specifically recognizes core snRNPs. Based on these findings, we propose that coilin serves as a quality control factor that selectively recognizes and sequesters incomplete snRNPs in CBs.

## Results

### Coilin specifically interacts with immature snRNPs

It has been previously shown that the C-terminal domain of coilin encompassing 214 amino acids (C214) associates with snRNPs, but the maturation status of these complexes has not been tested ^57,64^. To identify the fraction of snRNPs that interact with coilin, we incubated HeLa^coilinKO^ nuclear extract with recombinant GST-C214, pulled down the associated complexes, and resolved them by ultracentrifugation in a glycerol gradient (Fig. 1A). In all cases, snRNPs that associated with GST-C214 sedimented more slowly than snRNP particles in the nuclear extract, indicating that coilin preferentially interacts with incomplete snRNPs. Specifically, we observed an enrichment of the 12S U2 particle in the GST-C214 pulldown (fractions 6-9), whereas the mature 17S particle was underrepresented in comparison to the nuclear extract run in parallel (fractions 11-12). Similarly, we did not detect any tri-snRNP complexes interacting with coilin, though the U4/U6•U5 tri-snRNP was readily detectable in fractions 14-17 of the nuclear extract. Coilin further pulled down the U4/U6 di-snRNP (fractions 8-11) and a small U5 snRNA-containing complex (fractions 6-8). The mature 20S U5 particle is much larger and sedimented in fractions 12-14. We therefore speculate that the small core U5 snRNP particle contains only U5 snRNA and Sm proteins (core U5), but we did not investigate the identity of this complex any further.

**Figure 1.**
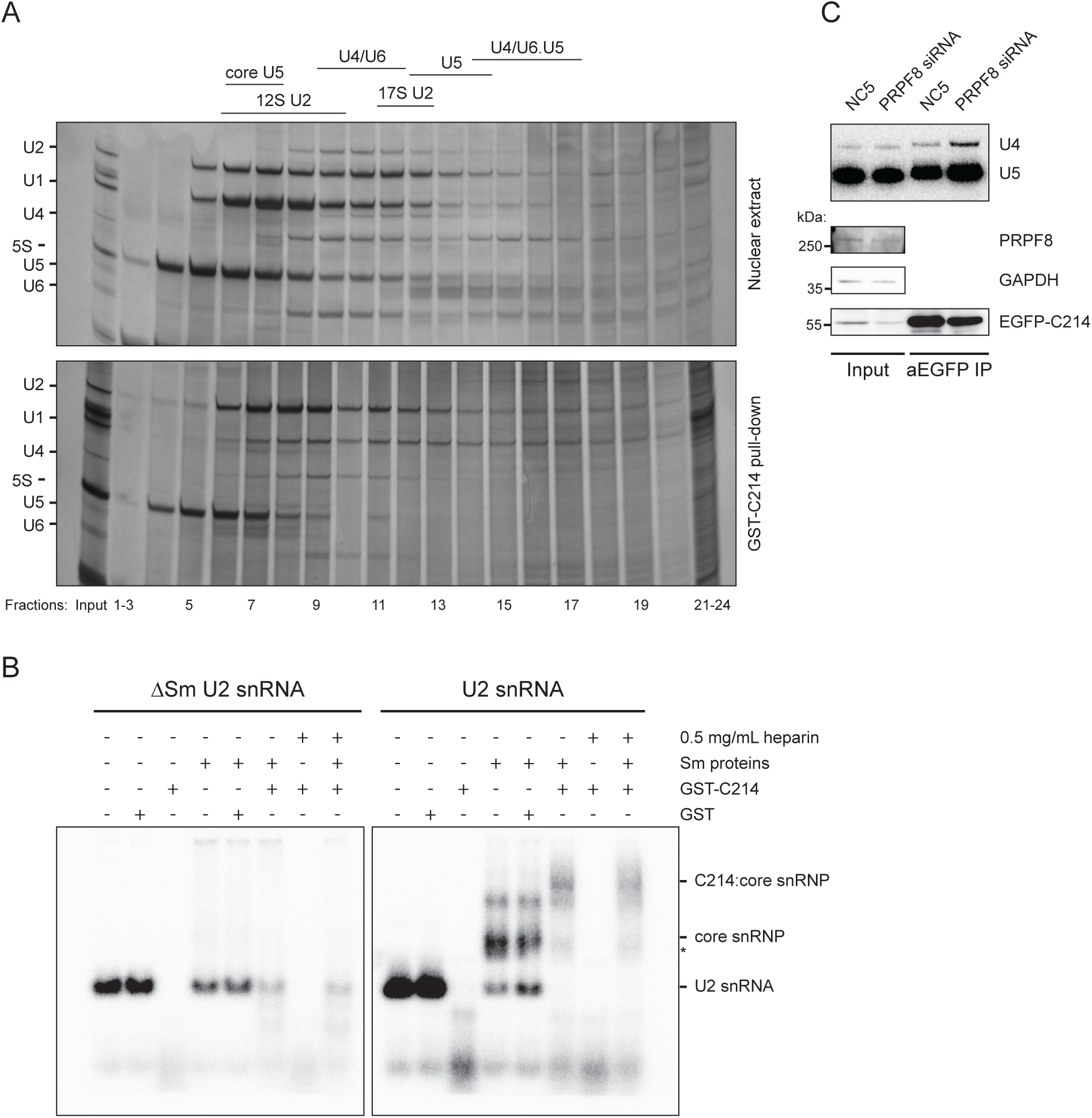
Coilin preferentially associates with immature snRNPs. A) SnRNP particles isolated from the nuclear extract (top) and particles associated with GST-C214 (bottom) were resolved using 10-30% glycerol gradient ultracentrifugation. SnRNAs in individual fractions were resolved on denaturing acrylamide RNA gels followed by silver-staining. Position of main snRNP particles indicated on top. B) Core U2-snRNP associates with GST-C214. Radioactively labeled U2 snRNA first interacts with Sm proteins forming the core snRNP. Core snRNP then associates with GST-C214 forming the C214:core snRNP complex. U2 lacking the Sm-binding site (ΔSm) served as a negative control. Complexes analyzed by the electrophoretic mobility shift assay. C) Downregulation of PRPF8 enhances association of U4 and U5 snRNAs with coilin EGFP-C214. snRNAs co-precipitated with transiently expressed EGFP-C214 were detected by Northern blotting.

To test whether coilin can directly interact with snRNPs, we assembled U2 core snRNP from *in vitro*-transcribed U2 snRNA and recombinant Sm proteins, and monitored the formation of complexes using electrophoretic mobility shift assay (EMSA) (Fig. 1B). GST-C214 associated with U2 core snRNP, showing that coilin directly associates with core snRNPs. In contrast, control reactions using U2 lacking the Sm site (ΔSm U2 snRNA) failed to form the core snRNP complex, and no shifts were observed with the addition of GST-C214 or GST alone. We observed substantial RNA degradation when snRNA was incubated with C214 without Sm proteins. We speculate that RNA was degraded by bacterial RNases co-purifying with recombinant GST-C214. Similarly, RNase activity has been previously observed in recombinant coilin isolated from bacteria ^56^. To further test whether coilin preferentially recognizes immature snRNPs, we knocked down PRPF8 to enrich the core U5 snRNPs and block tri-snRNP formation ^42^. PRPF8 depletion increased the association of U4 and U5 snRNAs with C214 (Fig. 1C), a finding that is consistent with our previous observation that PRPF8 downregulation enhances the sequestration of U4 and U5 snRNAs in CBs ^42^. Together, these data show that coilin directly interacts with core snRNPs preferentially in the context of incomplete assembly intermediates rather than mature splicing competent particles.

### Coilin C-terminus contains a bipartite snRNP interaction module

A previous study indicated that the interaction of the C-terminus of coilin with snRNPs is weakened in the absence of the RG box^57^. To test the contribution of the RG box in the interaction of coilin with snRNPs, we created a series of coilin deletions and/or mutations (Fig. 2A) and compared their binding with that of the wild-type (WT) protein. As a control, we include the K496E substitution in the Tudor-like domain, which has been shown to disrupt coilin-snRNP association ^46,64^. EGFP-tagged constructs were transiently expressed in Hela^coilinKO^ cells, immunoprecipitated by anti-EGFP antibodies, and coprecipitated snRNPs were monitored by detection of snRNP markers SmB and SNRPA1/U2A’ (Fig. 2B). Of all the constructs tested, the C214 fragment was the only one that recapitulated snRNP association observed with the full-length coilin, indicating that the C-terminal domain contains a minimal sequence sufficient for snRNP capture. Mutation or deletion of the RG box, as well as the K496E substitution, abolished coilin interaction with snRNPs, suggesting that both the RG box and the Tudor-like domain are necessary for productive snRNP binding. The RG box mutation had a similar effect when introduced in the context of C214 or full-length coilin, further confirming the importance of this motif for snRNP interaction. Next, we used unbiased mass spectrometry to assess the interaction network of the coilin C-terminal region. We prepared cell lysates from Hela^coilinKO^ cells, incubated them with recombinant WT GST-C214 or the inactive double mutant (R>A+K496E) and analyzed co-purified proteins by mass spectrometry (Fig. 2C). Interestingly, proteins of the arginine methylation machinery were the most differentially enriched hits, possibly hinting at a yet unknown regulatory network. Several snRNP-specific proteins, as well as all seven Sm proteins, were among the most differentially enriched hits for WT C214. This finding is consistent with the model that the RG-box and Tudor-like domains function together as a snRNP-binding module.

**Figure 2:**
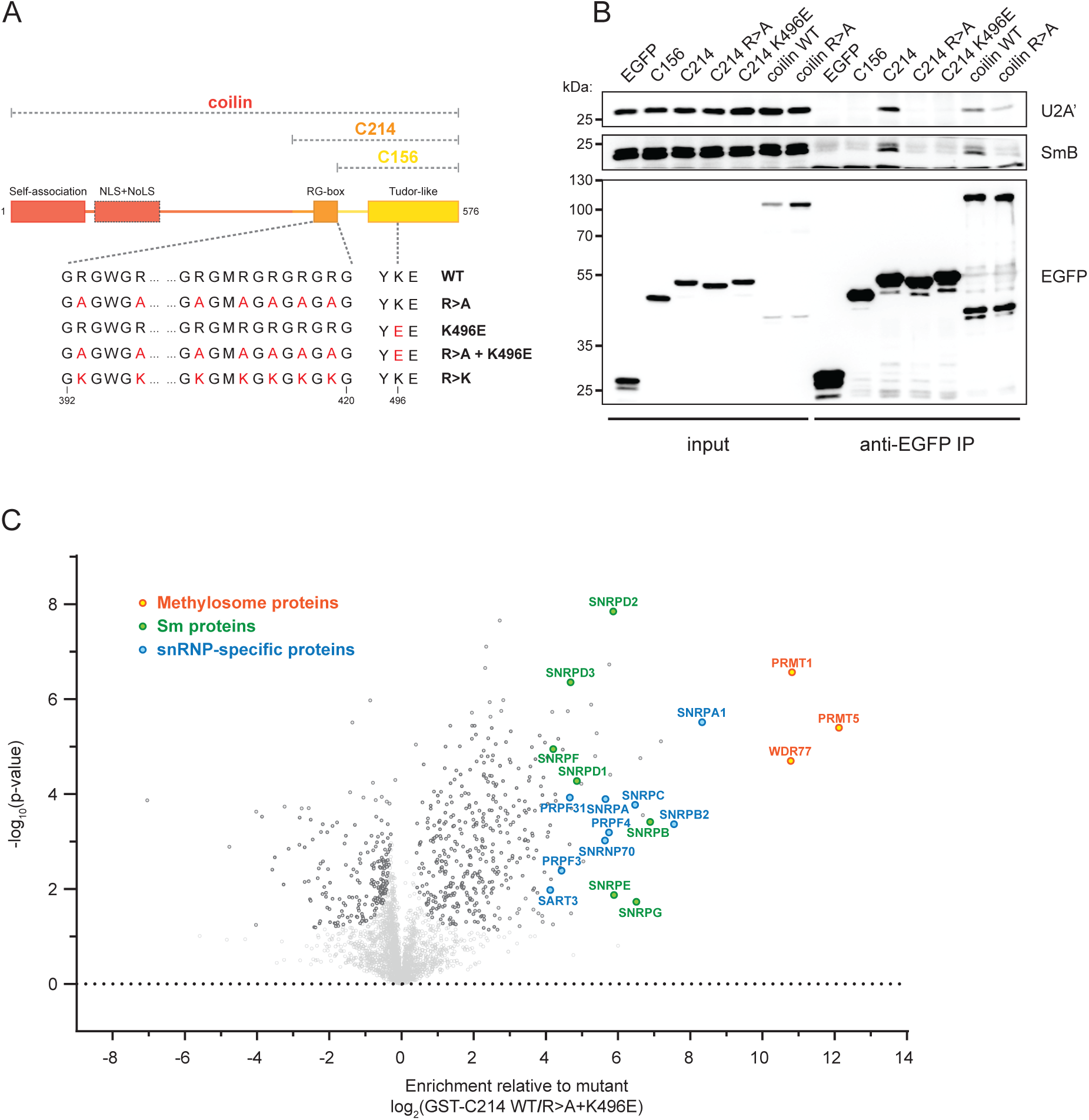
Coilin C-terminus interacts with snRNPs and methylation machinery. A) Schematic representation of various coilin mutants. Introduced substitutions marked in red. Position of several amino acids with respect to full-length coilin is indicated at the bottom. B) Mutations in the RG box and the Tudor-like domain impair coilin interaction with snRNPs. EGFP-C214 variants were transiently expressed in HeLa^coilinKO^ and immunoprecipitated. Co-precipitated snRNP markers U2A’ (U2 snRNP) and SmB protein (core snRNP) were detected by western blotting. C) A volcano plot depicting differential enrichment of proteins associated with WT GST-C214 relative to the double mutant (R>A+K496E) analyzed by mass spectrometry. The hits bellow the significance threshold are represented in light gray.

### Coilin interacts with RNA via the RG box

To further investigate the contribution of the RG box to snRNP binding, we prepared a series of R>A substitutions in order to identify key residues for the interaction. We observed a gradual loss of snRNP association correlating with an increased number of arginines substituted (Fig. S1). This finding suggests that the overall charge, rather than the exact position of individual arginines, is important for the interaction. To test this, we prepared a construct where we substituted all arginines in the RG box with lysines (C214 R>K) (Fig. 2A). We transiently expressed variants of EGFP-C214 in HeLa^coilinKO^ cells (Fig. 3A) or incubated cellular extracts from HeLa^coilinKO^ cells with recombinant variants of GST-C214 (Fig. 3B).

**Figure 3:**
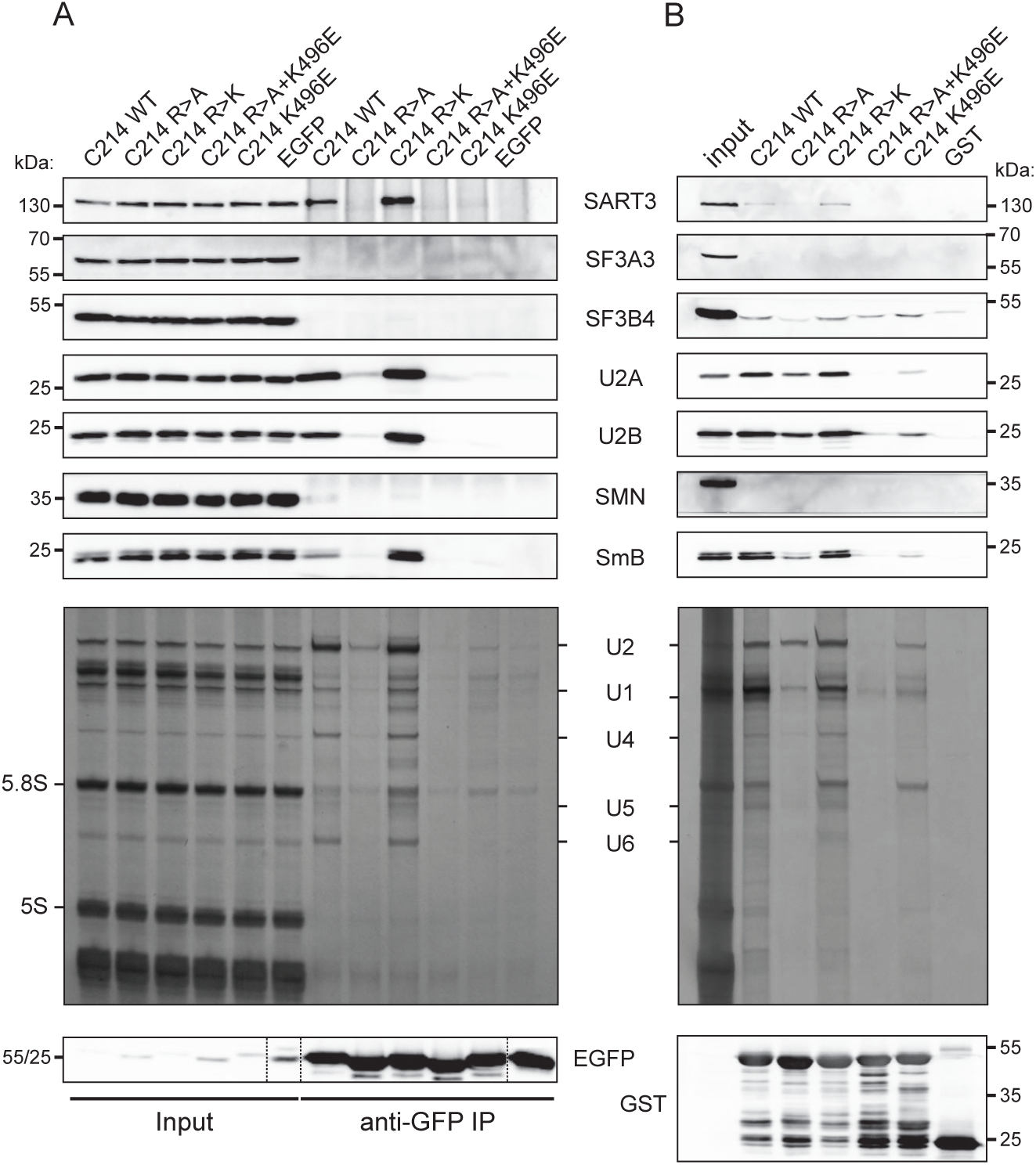
Both the RG box and the Tudor-like domain are necessary for snRNP interaction. A, B) Silver staining of RNAs and immunodetection of snRNP markers A) co-precipitated with transiently expressed EGFP-C214 variants and B) pulled down by recombinant GST-C214 variants.

Consistent with previous experiments, mutations in the RG box or the Tudor domain markedly reduced interactions with snRNAs and snRNP-specific proteins. The snRNP interaction was rescued by replacement of arginines by lysines (C214 R>K). These results demonstrate that the positively charged segment located upstream of the Tudor-like domain is critical for snRNP association and further confirm that both domains are necessary for productive snRNP binding. Both C214 WT and R>K mutant pulled down SART3, a marker of U4/U6 snRNP important for U4/U6 assembly and recycling ^22,27^ and SNRPA1/U2A’ and SNRPB2/U2B’’, which together interact with the U2 core snRNP to form the 12S assembly intermediate. In contrast, SF3B4 (15S U2 marker) and SF3A3 (17S U2 marker) were minimally interacting with C214. These findings are consistent with gradient ultracentrifugation (see Fig. 1A) and confirm that coilin interacts specifically with immature snRNPs. The SMN protein did not associate with recombinant GST-C214 lacking posttranslational modifications, which strongly indicates that the coilin-snRNP interaction is SMN-independent. These data are consistent with the model that coilin C-terminus is a snRNP-interacting module, which recognizes snRNPs via the synergic activity of the RG motif and the Tudor-like domain.

The necessity of a positive charge in the RG box hinted at a possible RNA interaction. To test whether C214 directly interacts with snRNAs, we incubated recombinant GST-C214 with *in vitro* transcribed snRNAs and observed strong interaction with all tested transcripts (Fig. 4A). Similarly to the EMSA (Fig. 1B), we observed RNA degradation when RNA was incubated with recombinant C214. However, due to the excess of RNA relative to protein compared to the stoichiometric RNA-protein ratio used in EMSA, we were able to detect RNA bound to C214. To determine whether the interaction is specific for snRNAs, we incubated GST-C214 with 7SK RNA, a highly structured short non-coding RNA that does not accumulate in CBs ^69^, and with MINX pre-mRNA ^70^. The C-terminal domain of coilin interacted with both 7SK and pre-mRNA to a similar extent as with U2 snRNA, indicating that the interaction of C214 with RNA is non-specific (Fig. 4B). To confirm that positively charged residues in the RG box facilitate the interaction with RNA, we tested the ability of RG box mutants to bind U2 snRNA (Fig. 4C). While replacement with alanines completely abolished the interaction, restoration of the positive charge by R>K rescued the RNA binding. As a control, we used the K496E substitution in the Tudor-like domain, which inhibits coilin association with snRNPs (Figs. 1-3 and ^64^), but this mutation did not affect C214 binding of U2 snRNA. These results show that the C-terminal domain interacts with various RNAs through the positively charged residues within the RG box, which form a non-specific RNA-binding domain.

**Figure 4:**
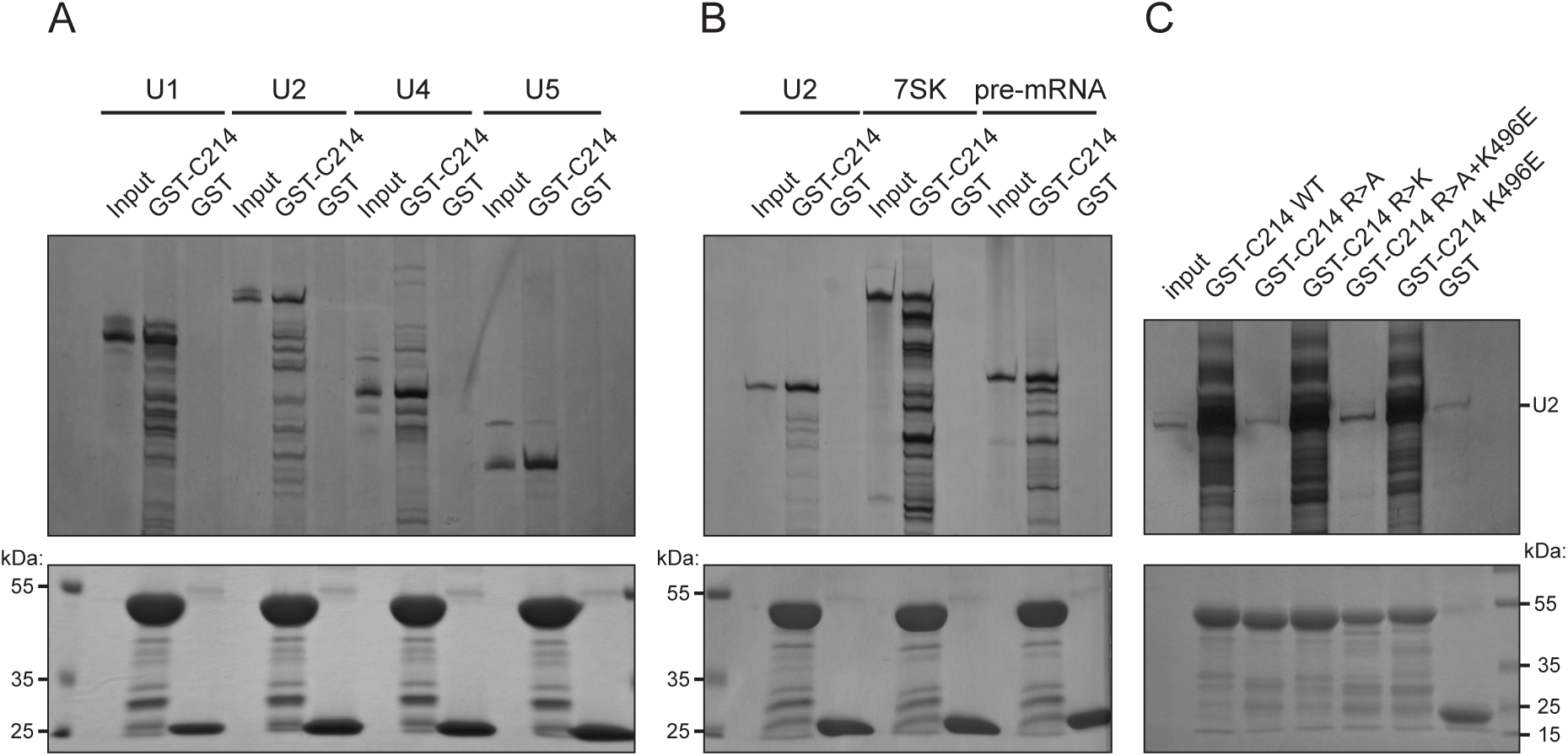
Coilin RG box is a novel RNA binding domain. *In vitro* transcribed RNAs were incubated with GST-C214 variants, resolved on denaturing RNA gels and visualized by silver staining (top). Coomassie staining of the captured recombinant proteins (bottom).

### Tudor-like domain recognizes SmE/F/G heterotrimerFEG

Next, we analyzed binding partner(s) of the Tudor-like domain. Previous studies have reported that this domain interacts with individual Sm proteins ^55,57,58^, but the exact binding partners and the molecular details of this interaction have remained unclear. We first modelled the structure of the Sm ring in complex with coilin Tudor-like domain consisting of 156 C-terminal amino acids by AlphaFold3 (Fig. 5A). The model predicted the heptameric ring of Sm proteins that closely resembles previously published experimental structures (Fig. S2) ^71,72^. Furthermore, the predicted structure of the coilin Tudor-like domain shares the key features with the previously published experimental structure, namely the β-barrel core and the extended loops ^61^. Interestingly, the model predicted the extended loops to be partially folded into antiparallel β-strands, giving them a more rigid conformation and contacting Sm proteins E, F and G (SNRPE, SNRPF and SNRPG).

**Figure 5:**
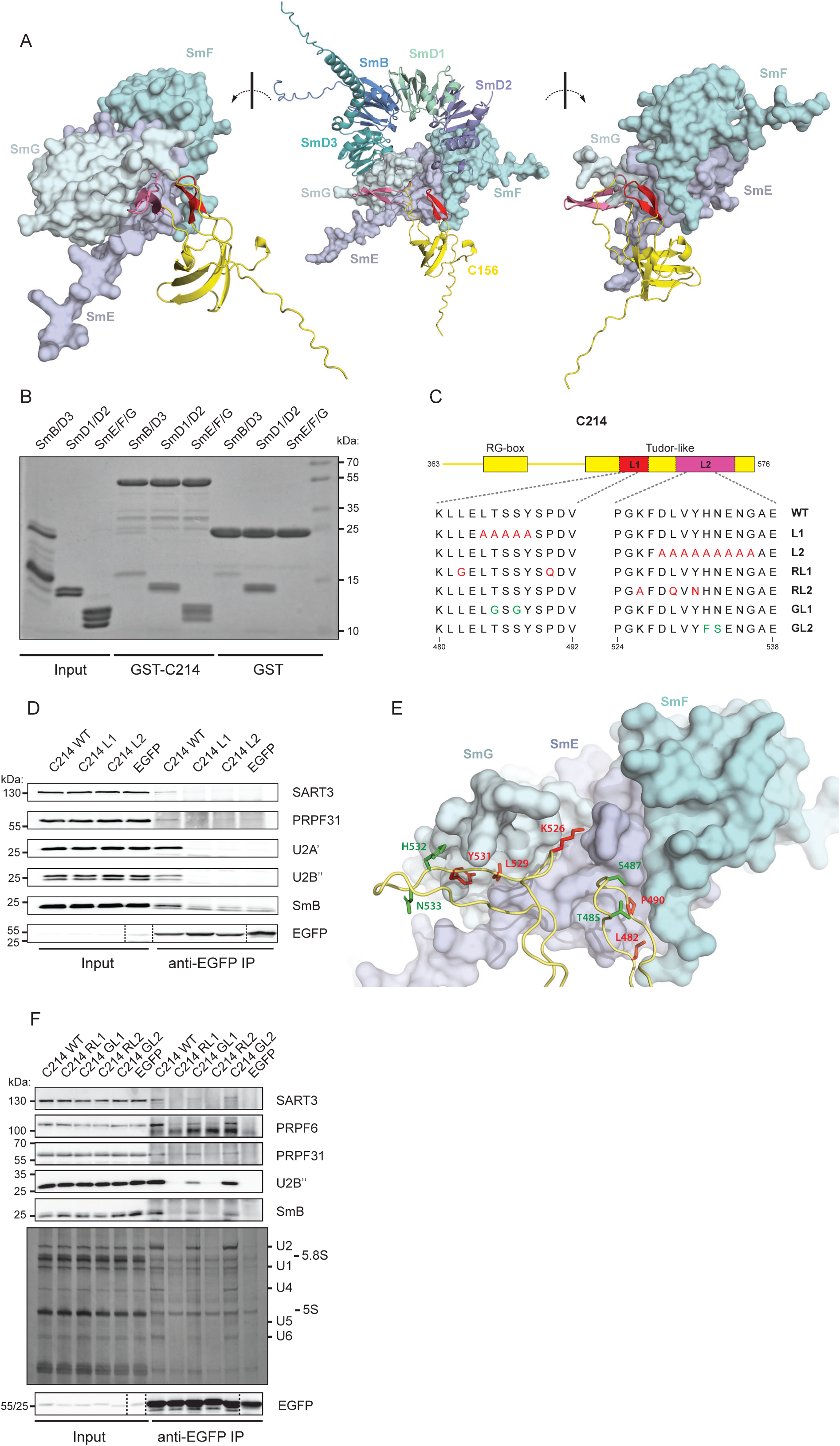
The Tudor-like domain interacts with SmE/F/G. A) AlphaFold 3 model of the C156-Sm ring complex. The two coilin loops interacting with the Sm proteins E, F and G are colored in red (β1-β2 loop/L1) and magenta (β3-β4 loop/L2). The unstructured N-terminus of C156 is removed to improve clarity. B) Recombinant C214 interacts specifically with the triplet E/F/G. Coomassie staining of the recombinant Sm complexes pulled down with recombinant GST-C214 or GST. Note the non-specific interaction of SmD1/D2 and SmB/D3 with GST alone. C) Schematic representation of the C214 variants with substitutions in loop1 (L1) and loop2 (L2). Position of several amino acids with respect to full-length coilin is indicated at the bottom. D) Immunodetection of snRNP markers co-precipitated with EGFP-C214 variants transiently expressed in HeLa^coilinKO^. E) A detailed view of the predicted interaction of loops 1 and 2 with the E/F/G triplet. Residues marked in red are potentially involved in the interaction with the Sm proteins, the residues marked in green are predicted not to be involved in the interaction. F) Substitution of amino acids in loops 1 and 2 predicted to interact with Sm proteins (RL1 and RL2) abolish C214-snRNP interaction. Substitution of amino acids in loops 1 and 2 predicted not to interact with Sm proteins (GL1 and GL2) maintain coilin-snRNP association. EGFP-C214 variants were transiently expressed in HeLa^coilinKO^ and co-precipitated proteins and RNAs detected by western blotting and silver staining, respectively.

To validate the AlphaFold3 model, we first incubated recombinant GST-C214 and GST-C156 with recombinant Sm subcomplexes B/D3, D1/D2 and E/F/G (Fig. 5B and Fig. S3). The trimeric E/F/G complex associated with both coilin constructs. The D1/D2 doublet, and to a lesser extent B/D3 doublet, non-specifically interacted with GST alone. However, we did not observe any increased binding to GST-C214 over GST alone, which indicates that these two dimers do not directly interact with coilin C-terminus. To examine the role of coilin loops in Sm ring binding, we introduced alanine substitutions within loop β1-β2 (C214 L1) or loop β3-β4 (C214 L2) (Fig. 5C). The alanine substitutions have been previously shown to preserve the structure of the Tudor-like domain ^61^. These two constructs were transiently expressed in HeLa^coilinKO^ cells and tested for snRNP association. Substitutions within either of the loops completely abolished the interaction with snRNPs, indicating that both loops are essential for efficient binding of Sm proteins (Fig. 5D). Next, we identified the coilin residues predicted by the AlphaFold3 model to interact with Sm proteins (Fig. 5E). We created several EGFP-C214 constructs mutating some of the potential Sm-interacting amino acids (marked in red). We also prepared two control constructs carrying the substitutions of the amino acids predicted not to interact with Sm proteins (marked in green). We transiently expressed these constructs in HeLa^coilinKO^ cells and performed immunoprecipitation using anti-EGFP antibodies (Fig. 5F). Mutations of amino acids predicted to bind Sm proteins abolished snRNP co-precipitation (C214 RL1 and RL2). The control constructs (C214 GL1 and GL2), on the other hand, maintained their association with snRNPs. Consistent with their importance, the interacting amino acids are evolutionary conserved across various species ^53^. These experiments provide an extensive validation of the predicted structure and offer explanation for the specificity of coilin-snRNP interaction.

The C-terminal region of coilin is a bipartite snRNP-interaction module recognizing core components of snRNPs - snRNA and Sm proteins - and both contacts are necessary to mediate the interaction between coilin and snRNPs. Individual bonds are apparently weak, as disruption of only one bond abolishes formation of the coilin-snRNP complex *in cellulo*, but together they provide a strong and specific platform for snRNP recognition and binding.

## Discussion

Coilin is a key scaffolding factor of the CB, but its molecular function has remained enigmatic since its discovery 35 years ago ^73,74^. Here, we characterized the C-terminal region of coilin and show that it contains two domains that together specifically recognize core snRNPs. The RG box interacts non-specifically with RNA, and the two conserved loops protruding from the Tudor core interact with three of the seven Sm proteins specific to snRNPs. This is consistent with previous findings that Sm proteins are critical for snRNP accumulation in the CBs ^43^.

Coilin has been shown to interact with RNA, but the exact position of the RNA-binding domain has not been identified. Earlier work placed the RNA-binding activity within the N-terminal region of coilin ^54,75,76^. Here, we show that the RG box located near the C-terminus binds RNA nonspecifically. This finding does not exclude that coilin contains additional RNA-binding domains, and RNA interaction with the coilin N-terminus has been suggested to be important for coilin multimerization and CB formation ^64^. The RG motif also interacts with SMN, and it is currently unclear whether SMN and RNA binding are mutually exclusive. SMN binding requires symmetric arginine methylation within the RG motif while the RNA interaction is not dependent on the methylation (Fig. 4) and it is currently unclear, whether RG methylation regulates RNA binding.

Tudor domains in many proteins recognize methylated arginine and lysine via an aromatic cage formed by four precisely positioned aromatic residues found in β1-β2 and β3-β4 loops. These loops are extended in the coilin Tudor-like domain, displacing the aromatic residues and disrupting the aromatic cage ^61^. Instead, the extended β1-β2 and β3-β4 loops have evolved into an alternative binding module that recognizes Sm proteins independently of the arginine methylation. The Tudor core itself is also important for snRNP interaction, because multiple mutations in the Tudor core abolish snRNP interaction ^64^. Based on the structure model (Fig. 5), we hypothesize that the Tudor core orients both loops in the position favorable for interaction with Sm proteins.

There is numerous evidence that CB sequesters immature and defective snRNPs ^16,17,42,43,77^. However, the CB factor(s) that specifically recognize immature particles have remained unidentified. Based on our finding that coilin C-terminal domain preferentially interacts with immature snRNPs (Figs. 1-3), we propose that coilin is the missing quality control factor that discriminates between mature and immature snRNPs and sequesters the immature particles in CB. The exact molecular mechanism of the release of mature snRNPs is currently unknown. We speculate that snRNP biogenesis and formation of the mature particles (the 17S U2 snRNP, 20S U5 and the U4/U6•U5 tri-snRNP) mask RNA and/or Sm core binding sites, thus weakening one of the binding interfaces and dissociating the entire complex (Fig. 6A). Consistently, superimposition of the predicted C214 - Sm ring structure onto the Sm ring of the 20S U5 snRNP ^78^ revealed a significant steric clash between EFTUD2 and the β3-β4 loop (L2) of the Tudor-like domain of coilin (Fig. 6B). Similarly, in the U4/U6•U5 tri-snRNP ^71^, the tri-snRNP specific protein SNRNP27 covers the surface of the SmE protein in U4 Sm ring, masking the β1-β2 loop (L1) binding interface predicted by AlphaFold3 (Fig. 6C). We did not find any potential clashes between coilin and snRNP-specific proteins in case of U1 and U2 snRNPs. In the case of U2 snRNP, however, we observe SF3A3 masking snRNA near their Sm ring, which might displace the RG box and thus weaken coilin-U2 association.

**Figure 6:**
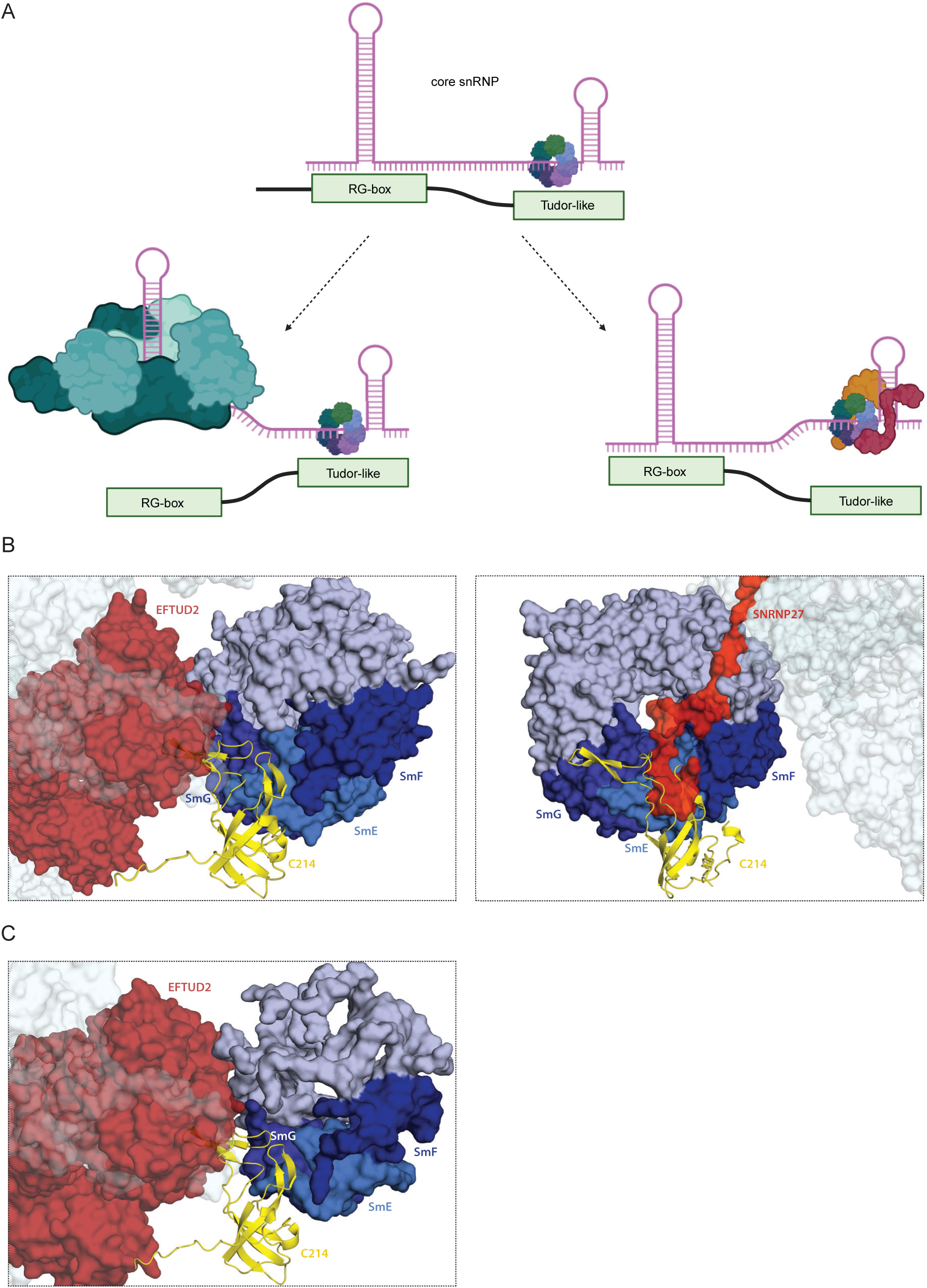
Model of coilin transient interaction with snRNPs during their biogenesis. A) Weakening of either RG box interaction with RNA or Tudor-like domain interaction with Sm proteins abolishes coilin-snRNP complex. B) Superimposition of AlphaFold 3 C214-Sm model onto U4 (left) and U5 Sm ring (right) of the U4/U6•U5 tri-snRNP. The tri-snRNP structure is based on PDB structure 6QW6 ^71^. Note the steric clash of tri-snRNP-specific protein SNRNP27 with coilin bound to U4 Sm ring and the U5 protein EFTUD2 with coilin bound to U5 Sm ring. Similar steric interference between coilin on U5 Sm ring and EFTUD2 is predicted using the structure of 20S U5 snRNP (PDB: 8RC0) ^78^.

Coilin was discovered 35 years ago as a marker of CBs, but the exact function of coilin remained enigmatic. Here, we provide evidence that coilin specifically interacts with immature snRNPs. Coilin oligomerization may thus bring various snRNPs to a close proximity, which provides molecular explanation for the proposed functions of CBs in promoting snRNP biogenesis. In addition, preferential binding of incomplete particles controls snRNP assembly and sequesters defective particles in CB.

## Material and Methods

### Cell culture

HeLa^coilinKO^ cell line ^46^ was cultured in high-glucose (4.5 g/L) DMEM (Sigma-Aldrich) supplemented with 10% FBS (Gibco) and 1% penicillin/streptomycin (Gibco) at 37 °C and 5% CO_2_.

### Plasmids

Other than C214 RL1 and RL2, which were obtained as dsDNA (GeneArt, ThermoFisher), all coilin constructs used in this manuscript are derived from EGFP-coilin^WT^ sequence ^46^. Coilin fragments (C214 and C156) were PCR-amplified using gene-specific primers and inserted into pEGFP-C1 plasmid using restriction enzyme cloning. Due to the loss of the endogenous NLS sequence (located near the N-terminus), TEV site and SV40 NLS sequences were added in frame, upstream of the gene of interest. Constructs carrying amino acid substitutions were prepared using site-directed mutagenesis and verified by DNA sequencing. For recombinant protein expression, selected constructs were subcloned into pGEX-6P-3 expression plasmid in-frame with GST (N-terminus) using restriction enzyme cloning. Plasmids for snRNA transcription were described previously ^43^.

**Antibodies**

|  |  |  |  |
| --- | --- | --- | --- |
| PRPF8 | ab79237 | 1:1000 | Abcam |
| GFP | sc-9996 | 1:500 | Santa Cruz |
| U2A | ab128937 | 1:4000 | Abcam |
| sDMA | Y12 | 1:4000 | produced in house from hybridoma cells |
| SMN | WH0006606M1 | 1:250 | Sigma-Aldrich |
| PRPF6 | sc-166889 | 1:500 | Santa Cruz |
| SART3 | ab176822 | 1:3000 | Abcam |
| SF3A3 | ab140605 | 1:4000 | Abcam |
| SF3B4 | ab183483 | 1:4000 | Abcam |
| U2B | TA808139 | 1:2000 | Thermo Fisher |
| GST | sc-138 | 1:1000 | Santa Cruz |
| PRPF31 | ab188577 | 1:4000 | Abcam |

### Transfection and transformation

pEGFP plasmids were transiently transfected into HeLa^coilinKO^ cells using Lipofectamine 3000 Transfection Reagent (Thermo Fisher Scientific) according to the manufacturer’s instructions. Cells were harvested 24h after transfection. pGEX plasmids were used to transform *E. coli* BL21-CodonPlus(DE3)-RIPL (Agilent) strain, selected on ampicillin (Amp) and chloramphenicol (Cam) supplemented agar plates. Individual positive colonies were cultured in Amp and Cam supplemented LB (LB + Amp + Cam) and stored as 25% glycerol stocks at - 80 °C.

### Protein purification

Overnight bacterial cultures grown in LB + Amp + Cam were used to inoculate fresh media to 0.2 OD. Fresh cultures were grown at 37 °C to 0.5 OD, followed by 1h at RT. Expression was then induced with 1 mM IPTG, and the cultures were cultivated at room temperature overnight. Cells were harvested by centrifugation and resuspended in lysis buffer (50 mM Tris-HCl pH7.5, 150 mM NaCl, 1 mM DTT) supplemented with lysozyme and phenylmethylsulfonyl fluoride (PMSF) (Fluka Biochemika), and lysed by sonication. Lysates were cleared by centrifugation, passed through 0.2 um filter and applied to a pre-equilibrated 1 mL GSTrap column (Cytiva). Bound proteins were washed with 10 column volumes (CV) of lysis buffer and eluted by the addition of 10 mM reduced glutathione. RNA-binding constructs (C214 WT, R>K, K496E) were diluted to 10 mM Tris-HCl pH7.5, 50 mM NaCl and applied to a pre-equilibrated 1 mL HiTrap Heparin column. Bound proteins were washed with 5 CV wash buffer (10 mM Tris-HCl pH7.5, 50 mM NaCl) and eluted by a linear salt gradient (50 mM -1 M). Fractions containing the protein of interest (∼500-600 mM NaCl) were measured on NanoDrop for nucleic acid contamination (A260/280<0.6). Following PAGE analysis, selected fractions were concentrated using Amicon 10 kDa cutoff ultrafiltration devices, aliquoted and snap frozen, and stored at -80 °C. Recombinant Sm proteins were produced and purified as described previously ^5,79^.

### Mass spectroscopy

3xGST pull-down - 20 ug of recombinant WT/R>A+K496E protein bound to 20 μL GST beads, incubated with HeLa^coilinKO^ cell lysates overnight, washed 4x with 50 mM Tris pH 7.5, 150 mM NaCl. All washing buffer aspirated, beads snap-frozen and stored at -80 °C. Beads were processed following the SP4 plus glass beads protocol ^80^. Briefly, samples were solubilized in 2% SDS prepared in 100 mM triethylammonium bicarbonate (TEAB) buffer, reduced with 10 mM tris(2-carboxyethyl)phosphine (TCEP), and alkylated with 40 mM chloroacetamide (performed together at 95 °C for 10 min). Proteins were digested overnight at 37 °C with MS-grade Lys-C (Wako, #125-05061) and trypsin (Promega, Cat# V5280) at a 1:40 enzyme-to-protein ratio. The resulting peptides were desalted using C18 StageTips and approx. 500 ng of peptide digest was separated on a Aurora Ultimate TS 25 cm x 75 µm C18 column using 90 min elution gradient on nanoUHPLC (Dionex Ultimate 3000) and analyzed in a DDA mode on an Orbitrap Exploris 480 mass spectrometer equipped with a FAIMS unit set to CV -40 and - 60 V. DDA MS Thermo raw files were converted to mzXML files each containing separate compensation voltage (CV, -40, -60 V) scans using FAIMS MzXML Generator. MzXML files were analyzed in MaxQuant (v. 2.6.6.0) with label-free quantification (LFQ) algorithm MaxLFQ and match between runs feature activated. FDR was set as 0.01 at all levels. UniProt human fasta proteome file was used (UP000005640_9606.fasta, UniProt Release 2024_01). Downstream processing of the proteinGroups.txt file was performed in Perseus (v2.1.3.0) using LFQ intensity values.

### RNAi

Custom Silencer® Select (Ambion) siRNA against PRPF8 (5’>3’ CCUGUAUGCCUGACCGUUtt/ AAACGGUCAGGCAUACAGGtt) were transfected using the Oligofectamine Transfection Reagent (Thermo Fisher Scientific) according to the manufacturer’s protocol. A single transfection was performed to freshly plated HeLa^coilinKO^ cells to a final concentration of 50 nM. Cells were then transfected with pEGFP-C214 construct 24h post-siRNA transfection and harvested the following day.

### Northern blot

RNA samples were resolved on 7 M urea denaturing polyacrylamide gel and transferred to Zeta-Probe membrane (Bio-Rad) using semi-dry method. The transfer was performed for 1 h in 0.5xTBE buffer at 15 V, and transfered RNA was UV crosslinked to the membrane by 120 mJ/cm2. Membrane was then pre-hybridized in SES1 buffer (0.5 M Sodium phosphate pH7.2, 7% SDS, 10 mM EDTA) for 30 min at 37 °C. The DNA oligonucleotide probes complementary to U5 and U4 snRNA were terminally labeled with [γ-^32^P]ATP (Hartman Analytic) using T4 polynucleotide kinase, purified on Micro-Spin G25 columns (GE Healthcare) and denatured at 95 °C for 5 min. Probes were added to SES1 buffer and hybridized on the membrane at 37 °C overnight. Membrane was then washed 2x 20 min at 37 °C with 1xSSC (15 mM Sodium citrate pH7, 150 mM NaCl) suplemented with 0.1% SDS, exposed to Storage Phosphor Screen overnight, and scanned on Amersham Typhoon 9500 (GE Healthcare).

### RNA *in vitro* transcription

snRNA and 7SK RNA templates were amplified by PCR using forward primer containing T7 promoter sequence and gene-specific reverse primer. MINX pre-mRNA template was prepared by linearization of plasmid containing MINX sequence under the control of T7 promoter. Transcription reactions were assembled with MEGAshortscript™ T7 transcription kit (Thermo Fisher) and incubated for 4h at 37 °C. RNA was isolated by phenol/chloroform extraction and dissolved in nuclease-free water. Radiolabeled RNA was prepared by addition of [α-^32^P]UTP (Hartman Analytic) in the transcription reaction and isolated using Direct-zol™ RNA Miniprep Kit.

### Electrophoretic mobility shift assay

SmE/F/G, SmD1/D2, and SmD3/B were diluted in Sm buffer to 4 µM final concentration. For snRNP core assembly with GST-C214, 8 pmol of preassembled Sm proteins were incubated with 8 pmol radiolabeled U2 snRNA in Sm buffer supplemented with human RNase inhibitor (Diana Biotechnologies), 10 μg/ml BSA, and 10 μg/ml yeast RNA for 1h at 37 °C. After incubation, 8 pmol of GST-C214 or GST were added, and reactions incubated for 30 min at 4 °C. Assembled complexes were mixed with 5x loading dye with or without heparin (0.5 mg/ml) and analyzed on 6% native PAGE gel (37.5:1) run in 0.5xTBE buffer. The gel was exposed to a phosphor screen for 24-48 hours and scanned on a phosphor imager (Typhoon^TM^, Amersham). Radiolabeled U2 snRNA lacking the Sm-site sequence (AUUUUUG, ΔSm U2 snRNA) was prepared along with the WT sequence of U2 snRNA and used as a negative control.

### Immunoprecipitation and pull-down

Cells were harvested by trypsinization at confluency and resuspended in NET2 buffer (50 mM Tris-HCl pH 7.4, 150 mM NaCl, 0.05% Nonidet P-40) supplemented with Protease Inhibitor Cocktail Set III (Millipore) and RNasin ribonuclease inhibitor (Promega). After 2x45 s sonication (0.5 s at 50% of maximum energy), the lysates were centrifuged at 20 000 × g, 4 °C for 10 min, and the supernatants precleared with agarose beads for 1 h at 4 °C. Three percent of the lysates were saved for input control, while the rest was incubated with anti-EGFP antibodies coupled to Protein G PLUS agarose beads overnight at 4 °C. Bound complexes were washed with NET2 buffer and solubilized in TRIzol® Reagent (Ambion). RNA was extracted using phenol/chloroform, dissolved in urea sample buffer (20 mM Tris-HCl pH 8.0, 8 M urea, 0.2% xylene blue), resolved on a 7 M urea denaturing polyacrylamide gel, and analyzed by silver staining or northern blotting. Proteins were precipitated with isopropanol, dissolved in SDS sample buffer (0.25 M Tris-HCl pH 6.8, 20% glycerol, 4% SDS, 2% β-mercaptoethanol, 0.02% bromphenol blue), resolved on a 12% polyacrylamide gel, and analyzed by Coomassie staining or western blotting. For pull-down assays, 50 ug recombinant GST-tagged protein was bound to Pierce Glutathione Agarose beads (Thermo Fisher), washed with NET2 buffer and incubated with cell lysates, in vitro transcribed RNA, or recombinant Sm proteins. Pull-down of Sm proteins was performed in Sm buffer (20 mM HEPES-NaOH pH 7.5, 200 mM NaCl, 5 mM DTT) instead of NET2.

### Glycerol-gradient centrifugation

Pierce Glutathione Agarose beads were incubated with 100 ug recombinant GST-C214, washed with NET2 buffer, and incubated with HeLa^coilinKO^ lysates overnight at 4 °C. Bound complexes were washed with NET2 buffer and eluted with 10 mM reduced GSH. Nuclear extracts were prepared from HeLa^coilinKO^ using NE-PER nuclear and cytoplasmic extraction reagents (Thermo Fisher Scientific) according to the manufacturer’s instructions. Samples were diluted in gradient buffer (20 mM HEPES/KOH pH 8.0, 150 mM KCl, 1.5 mM MgCl 2) supplemented with Protease Inhibitor Cocktail Set III (Millipore), 0.5 mM PMSF, and 0.5 mM DTT and loaded on a linear 10–30% glycerol gradient. Complexes were centrifuged at 130,000×g, 4 °C for 17h and fractionated (24 fractions, 500 μL each). RNA was extracted from each fraction using phenol/chloroform, resolved on 7 M urea denaturing polyacrylamide gel and analyzed by silver staining.

## Acknowledgement

The author thanks Dr. Marek Vrbacky, Poulami Banik, Jana Wiedermanova and Jana Machatova-Krizova for technical assistance. This work was supported by grants from the Czech Science Foundation (24-11157S), the Project JAC CZ.02.01.01/00/22_008/0004575 “RNA for therapy”, co-funded by the European Union and Grant Agency of Charles University (No. 264123 to N.R.)

## Author contribution

D.S., M.G and N.R. conceived the project; N.R., M.G. and V.H. performed the experiments; U.F. provided material and protocols; D.S. and N.R. manuscript writing and funding.

## Supplementary figure legend

**Figure S1:**
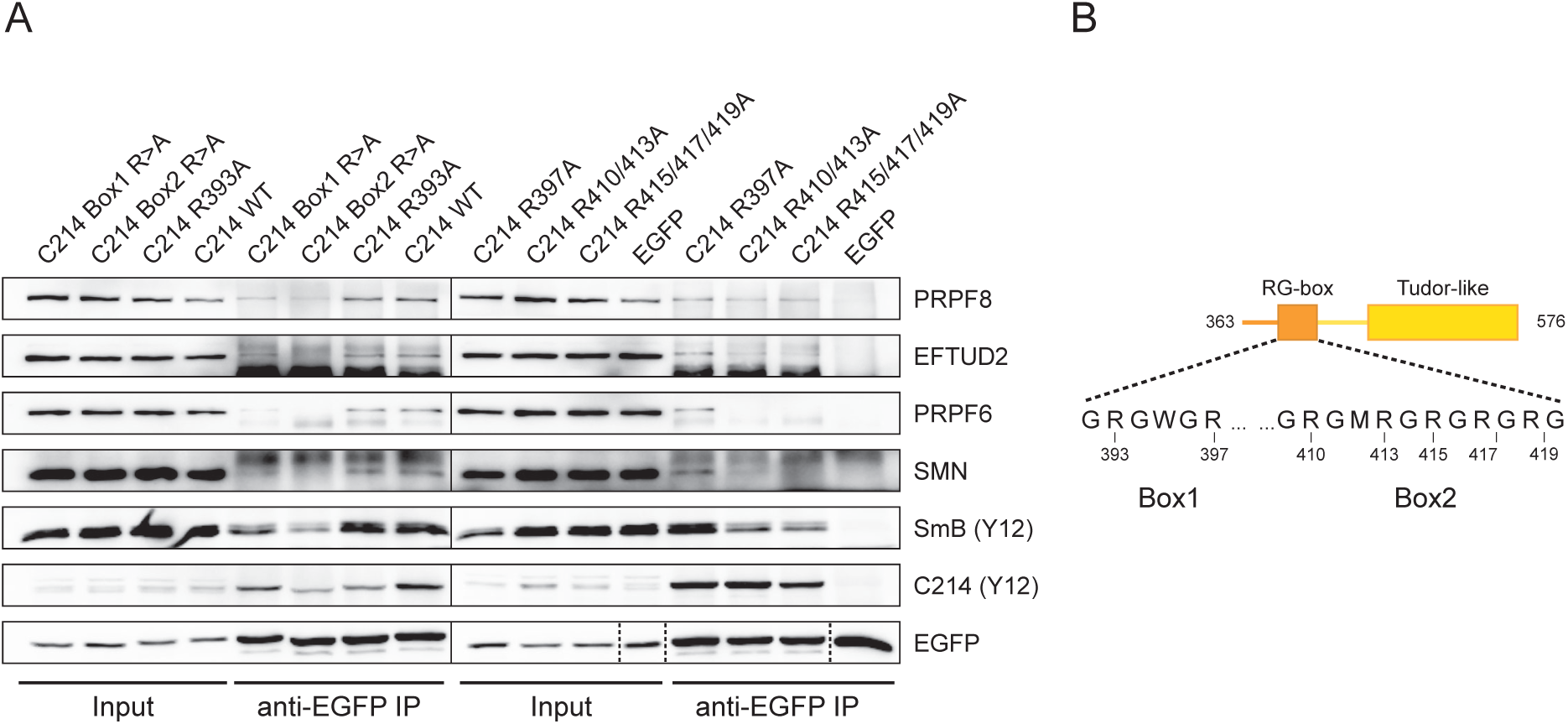
Contribution of individual arginines within the RG box towards snRNP binding. A) Immunodetection of different snRNP markers and symmetrical dimethylarginines (Y12 antibody) following a co-immunoprecipitation with transiently expressed EGFP-C214 RG box variants. Samples were resolved on two separate gels. B) Schematic representation of C214 with the sequence of the two arginine-rich segments (Box1 and Box2) of the RG box denoting the positions of mutated residues.

**Figure S2:**
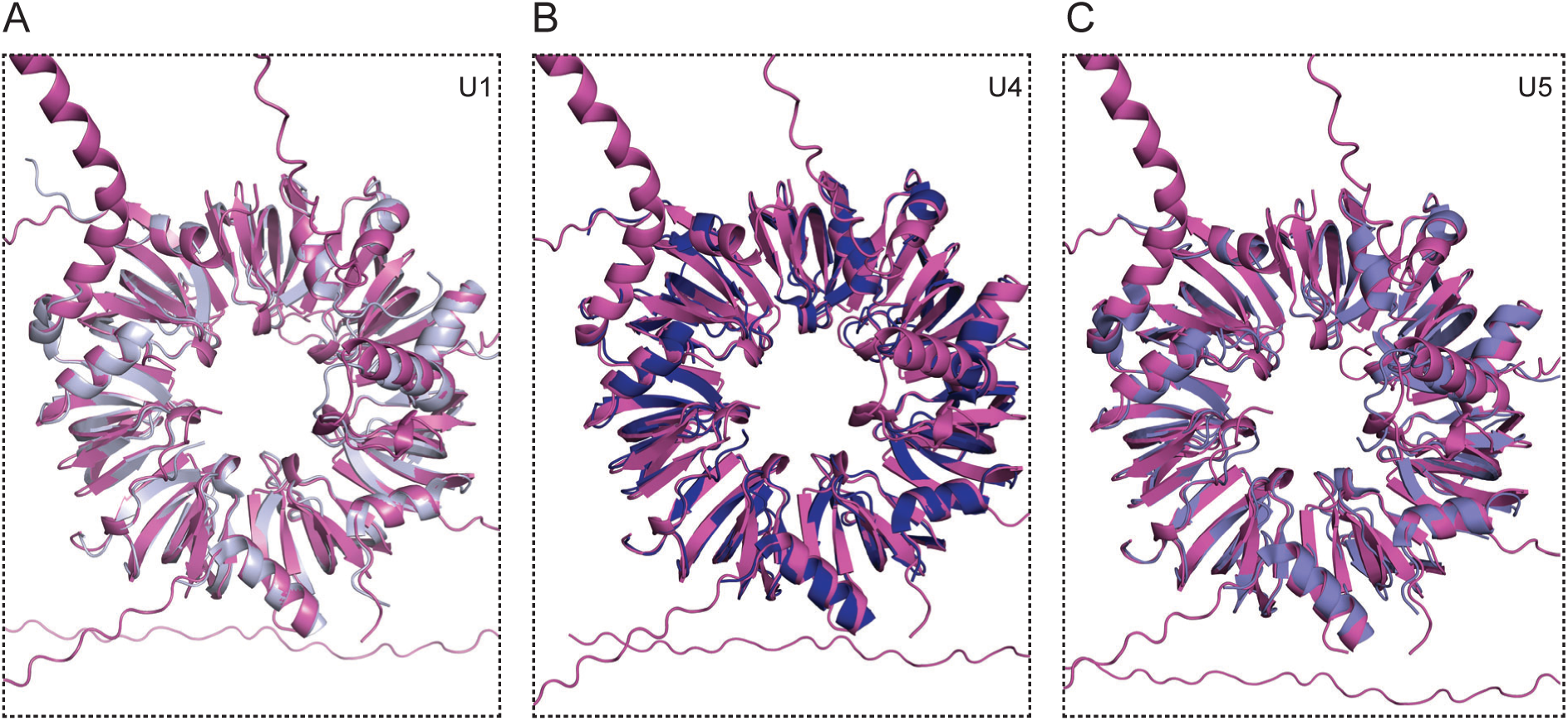
Comparison of the predicted and experimental Sm ring structures. Superimposition of AlphaFold 3 predicted structure of the Sm ring onto experimentally determined structure of A) U1 Sm ring (PDB: 3CW1), B) U4 Sm ring (PDB: 6QW6) and C) U5 Sm ring (PDB: 6QW6)

**Figure S3:**
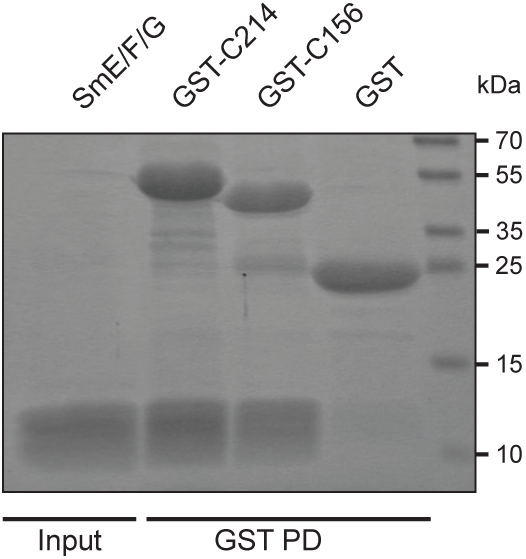
Recombinant GST-C156 interacts with Sm E/F/G triplet. Coomassie staining of proteins resolved on acrylamide gel following a GST pull-down.

**Figure S4:**
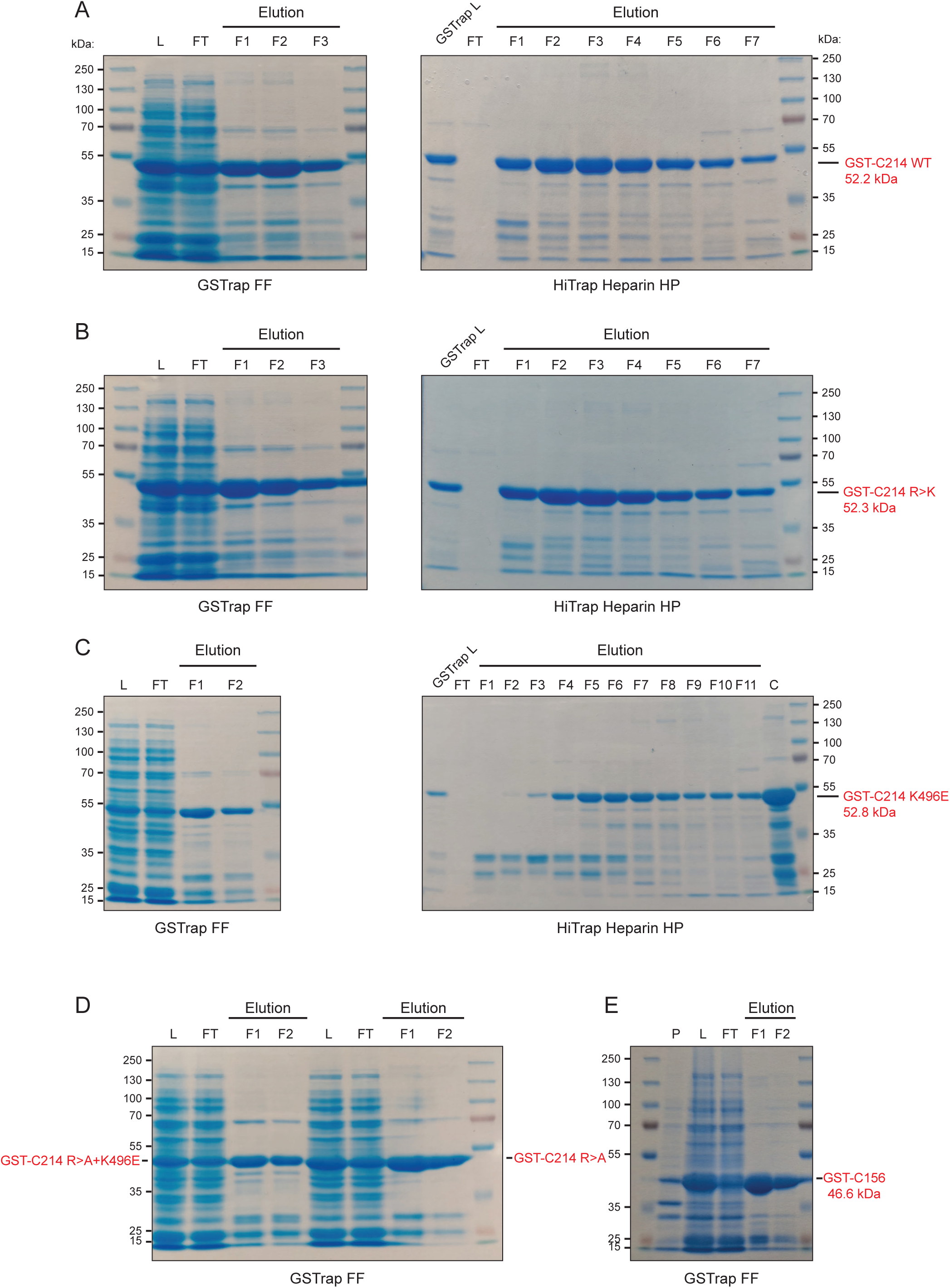
Purification of recombinant GST-tagged coilin variants and mutants. A) GST-C214 WT, B) GST-C214 R>K, C) GST-C214 K496E, D) GST-C214 R>A and GST-C214 R>A+K496E, E) GST-C165. Legend: L - load/clarified bacterial lysate; FT - flow-through; F# - fraction number; C - concentrated protein elution; P - pellet.

